# Whole-genome sequencing is more powerful than whole-exome sequencing for detecting exome variants

**DOI:** 10.1101/010363

**Authors:** Aziz Belkadi, Alexandre Bolze, Yuval Itan, Aurélie Cobat, Quentin B. Vincent, Alexander Antipenko, Lei Shang, Bertrand Boisson, Jean-Laurent Casanova, Laurent Abel

## Abstract

We compared whole-exome sequencing (WES) and whole-genome sequencing (WGS) in six unrelated individuals. In the regions targeted by WES capture (81.5% of the consensus coding genome), the mean numbers of single-nucleotide variants (SNVs) and small insertions/deletions (indels) detected per sample were 84,192 and 13,325, respectively, for WES, and 84,968 and 12,702, respectively, for WGS. For both SNVs and indels, the distributions of coverage depth, genotype quality, and minor read ratio were more uniform for WGS than for WES. After filtering, a mean of 74,398 (95.3%) high-quality (HQ) SNVs and 9,033 (70.6%) HQ indels were called by both platforms. A mean of 105 coding HQ SNVs and 32 indels were identified exclusively by WES, whereas 692 HQ SNVs and 105 indels were identified exclusively by WGS. We Sanger sequenced a random selection of these exclusive variants. For SNVs, the proportion of false-positive variants was higher for WES (78%) than for WGS (17%). The estimated mean number of real coding SNVs (656, ∼3% of all coding HQ SNVs) identified by WGS and missed by WES was greater than the number of SNVs identified by WES and missed by WGS (26). For indels, the proportions of false-positive variants were similar for WES (44%) and WGS (46%). Finally, WES was not reliable for the detection of copy number variations, almost all of which extended beyond the targeted regions. Although currently more expensive, WGS is more powerful than WES for detecting potential disease-causing mutations within WES regions, particularly those due to SNVs.

**Significance:** Whole-exome sequencing (WES) is gradually being optimized to identify mutations in increasing proportions of the protein-coding exome, but whole-genome sequencing (WGS) is becoming an attractive alternative. WGS is currently more expensive than WES, but its cost should decrease more rapidly than that of WES. We compared WES and WGS on six unrelated individuals. The distribution of quality parameters for single-nucleotide variants (SNVs) and insertions/deletions (indels) was more uniform for WGS than for WES. The vast majority of SNVs and indels were identified by both techniques, but an estimated 650 high-quality coding SNVs (∼3% of coding variants) were detected by WGS and missed by WES. WGS is therefore slightly more efficient than WES for detecting mutations in the targeted exome.

## Introduction

Whole-exome sequencing (WES) is routinely used and is gradually being optimized for the detection of rare and common genetic variants in humans (1–8). However, whole-genome sequencing (WGS) is becoming increasingly attractive as an alternative, due to its broader coverage and decreasing cost (9–11). It remains difficult to interpret variants lying outside the protein-coding regions of the genome. Diagnostic and research laboratories, whether public or private, therefore tend to search for coding variants, most of which can be detected by WES, first. Such variants can also be detected by WGS, and several studies previously compared WES and WGS for different types of variations and/or in different contexts (9, 11–16), but none of them in a really comprehensive manner. Here, we compared WES and WGS, in terms of detection rates and quality, for single-nucleotide variants (SNVs), small insertions/deletions (indels), and copy number variants (CNVs) within the regions of the human genome covered by WES, using the most recent next-generation sequencing (NGS) technologies. We aimed to identify the most efficient and reliable approach for identifying these variants in coding regions of the genome, to define the optimal analytical filters for decreasing the frequency of false-positive variants, and to characterize the genes that were either hard to sequence by either approach or were poorly covered by WES kits.

## Results

We compared the two NGS techniques, by performing WES with the Agilent Sure Select Human All Exon kit 71Mb (v4 + UTR), and WGS with the Illumina TruSeq DNA PCR-Free sample preparation kit on blood samples from six unrelated Caucasian patients with isolated congenital asplenia (OMIM #271400) (6). We used the genome analysis toolkit (GATK) best-practice pipeline for the analysis of our data (17). We used the GATK Unified Genotyper (18) to call variants, and we restricted the calling process to the regions covered by the Sure Select Human All Exon kit 71Mb plus 50 base pairs (bps) of flanking sequences on either side of each of the captured regions, for both WES and WGS samples. These regions, referred to as the WES71+50 region, included 180,830 full-length and 129,946 partial exons from 20,229 protein-coding genes (**Table 1**). There were 65 million reads per sample, on average, mapping to this region in WES, corresponding to a mean coverage of 73X (**Table S1**), consistent with the standards set by recent large-scale genomic projects aiming to decipher disease-causing variants by WES (9, 22). On average, 35 million reads per sample mapped to this region by WGS, corresponding to a mean coverage of 39X (**Table S1**). We first focused on the analysis of single-nucleotide variants (SNVs). The mean (range) number of SNVs detected was 84,192 (82,940-87,304) by WES and 84,968 (83,340-88,059) by WGS. The mean number of SNVs per sample called by both methods was 81,192 (∼96% of all variants) (**Fig. S1A)**. For 99.2% of these SNVs, WES and WGS yielded the same genotype, and 62.4% of these concordant SNVs were identified as heterozygous (**Fig. S1B**). These results are similar to those obtained in previous WES studies (22, 1, 5). Most of the remaining SNVs (329 of 415) with discordant genotypes for these two techniques were identified as homozygous variants by WES and as heterozygous variants by WGS (**Fig. S1B**).

**Table 1:** Specific regions of the genome covered by WES using the 71Mb kit. Four types of genomic units were analyzed: exons from protein-coding genes, miRNA exons, snoRNA exons, and lincRNA exons as defined in Ensembl Biomart (19). We determined the number of these units using the R Biomart package (20) on the GRCh37/hg19 reference. We first considered all exons from protein-coding genes (denoted as All) obtained from Ensembl. The essential splice sites (i.e. the two intronic base pairs at the intron/exon junction) were not included in our analysis of exons. Then we focused on protein-coding exons with a known CDNA coding start and CDNA coding end, and present in CCDS transcripts (21). For the counts, we excluded one of the duplicated units of the same type, or units entirely included in other units of the same type (only the longest unit would be counted in this case). We then determined the number of the remaining units that were fully or partly covered when considering the genomic regions defined by the Agilent Sure Select Human All Exon kit 71Mb (v4 + UTR) with the 50 bps flanking regions.

|  | Exons from protein-coding genes |  | lincRNA | miRNA | snoRNA |
| --- | --- | --- | --- | --- | --- |
|  | All | CCDS |  |  |  |
|  | 71Mb +/- 50Bps | 71Mb +/- 50Bps | 71Mb +/- 50Bps | 71Mb +/- 50Bps | 71Mb +/- 50Bps |
| Fully included | 180,830 | 147,131 | 554 | 1,171 | 252 |
| Partially included | 129,946 | 34,892 | 855 | 94 | 93 |
| Fully excluded | 64,921 | 5,762 | 25,389 | 1,782 | 1,111 |
| Total | 375,697 | 187,785 | 26,798 | 3,047 | 1,456 |

We then investigated, in WES and WGS data, the distribution of the two main parameters assessing SNV quality generated by the GATK variant calling process (18): coverage depth (CD), corresponding to the number of aligned reads covering a single position; and genotype quality (GQ), which ranges from 0 to 100 (higher values reflect more accurate genotype calls). We also assessed the minor read ratio (MRR), which was defined as the ratio of reads for the less covered allele (reference or variant allele) over the total number of reads covering the position at which the variant was called. Overall, we noted reproducible differences in the distribution of these three parameters between WES and WGS. The distribution of CD was skewed to the right in the WES data, with a median at 50X but a mode at 18X, indicating low levels of coverage for a substantial proportion of variants (**Fig. 1A**). By contrast, the distribution of CD was normal-like for the WGS data, with the mode and median coinciding at 38X (**Fig. 1A**). We found that 4.3% of the WES variants had a CD < 8X, versus only 0.4% of the WGS variants. The vast majority of variants called by WES or WGS had a GQ close to 100. However, the proportion of variants called by WES with a GQ < 20 (3.1%) was, on average, twice that for WGS (1.3%) (**Fig. 1B**). MRR followed a similar overall distribution for WES and WGS heterozygous variants, but peaks corresponding to values of MRR of 1/7, 1/6, 1/5 and 1/4 were detected only for the WES variants (**Fig. 1C**). These peaks probably corresponded mostly to variants called at a position covered by only 7, 6, 5 and 4 reads, respectively. The overall distributions of these parameters indicated that the variants detected by WGS were of higher and more uniform quality than those detected by WES.

**Fig. 1.**
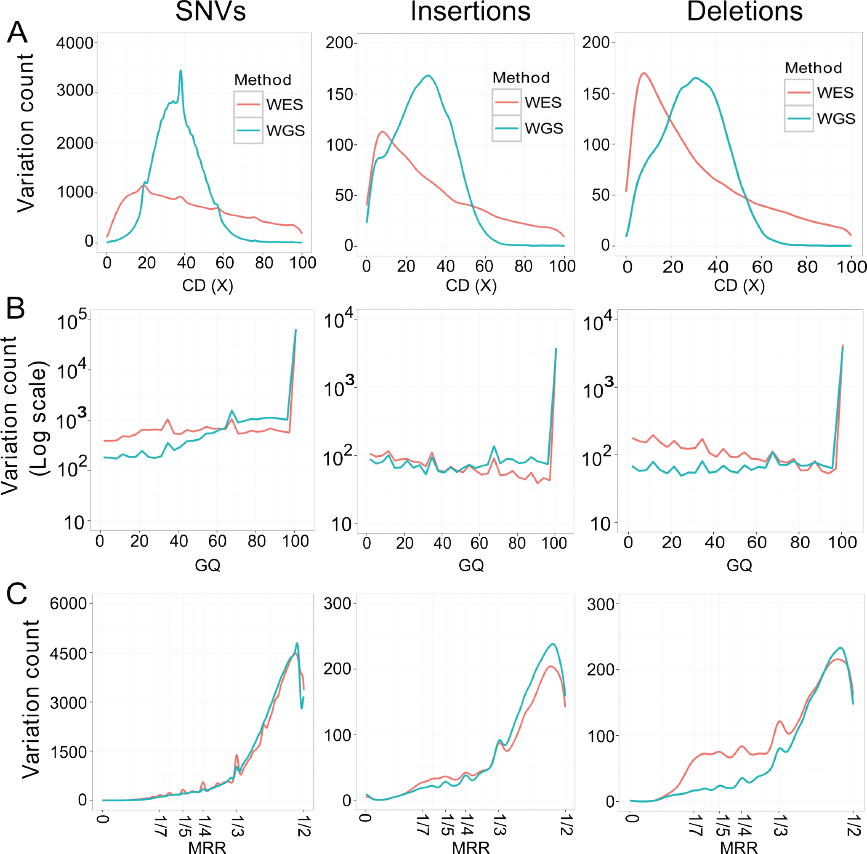
Distribution of the three main quality parameters for the variations detected by WES or WGS. **(A)** Coverage depth (CD), **(B)** genotype quality (GQ) score, and **(C)** minor read ratio (MRR). For each of the three parameters, we show the average over the 6 WES (red) and the 6 WGS (turquoise) samples in: SNVs (left panel), Insertions (middle panel) and Deletions (right panel).

Next, we looked specifically at the distribution of these parameters for the variants with genotypes discordant between WES and WGS, denoted as discordant variants. The distribution of CD for WES variants showed that most discordant variants had low coverage, at about 2X, with a CD distribution very different from that of concordant variants (**Fig. S2A**). Moreover, most discordant variants had a GQ < 20 and a MRR < 0.2 for WES (**Fig. S2B**). By contrast, the distributions of CD, GQ, and MRR were very similar between WGS variants discordant with WES results and WGS variants concordant with WES results (**Fig. S2**). All these results indicate that the discordance between the genotypes obtained by WES and WGS was largely due to the low quality of WES calls for the discordant variants. We therefore conducted subsequent analyses by filtering out low-quality variants. We retained SNVs with a CD ≥ 8X and a GQ ≥ 20, as previously suggested (24), and with a MRR ≥ 0.2. Overall, 93.8% of WES variants and 97.8% of WGS variants satisfied the filtering criterion (**Fig. S3A**). We recommend the use of these filters for projects requiring high-quality variants for analyses of WES data. More than half (57.7%) of the WES variants filtered out were present in the flanking 50 bp regions, whereas fewer (37.6%) of the WGS variants filtered out were present in these regions. In addition, 141 filtered WES variants and 70 filtered WGS variants per sample concerned the two bps adjacent to the exons, which are key positions for splicing. After filtering, the two platforms called an average of 76,195 total SNVs per sample, and the mean proportion of variants for which the same genotype was obtained with both techniques was 99.92% (range: 99.91%-99.93%).

We then studied the high-quality (HQ) variants satisfying the filtering criterion but called by only one platform. On average, 2,734 variants (range: 2,344-2,915) were called by WES but not by WGS (**Fig. S3A**), and 6,841 variants (5,623-7,231) were called by WGS but not WES (**Fig. S3A**). We used Annovar software (25) to annotate these HQ variants as coding variants, i.e., variants overlapping a coding exon, this term referring to the coding part of the exon but not the UTR portion. Overall, 651 of the 2,734 WES-exclusive HQ variants and 1,113 of the 6,841 WGS-exclusive HQ variants were coding variants (**Fig. S3A)**. Using the Integrative Genomics Viewer (IGV) tool (26), we noticed that most WES-exclusive HQ variants were also present on the WGS tracks with quality criteria that were above our defined thresholds. We were unable to determine why they were not called by the Unified Genotyper. We therefore used the GATK Haplotype Caller to repeat the calling of SNVs for the WES and WGS experiments. We combined the results obtained with Unified Genotyper and Haplotype Caller and limited subsequent analyses to the variants called by both callers. The mean number (range) of HQ coding SNVs called exclusively by WES fell to 105 (51-140) per sample, whereas the number called exclusively by WGS was 692 (506-802) (**Fig. S3B**) indicating that calling issues may account for ∼80% of initial WES exclusive coding variants and ∼40% of initial WGS exclusive coding variants. The use of variants identified by both Unified Genotyper and Haplotype Caller (i.e. the intersection) therefore appeared to increase the reliability and accuracy of calls, which served the main purpose of our study. Using variants identified by one or both callers only (i.e. the union), rather than the intersection, would increase the number of false positives but decrease the number of false negatives, and this might be of interest in specific contexts. Using the intersection, we obtained a mean of 74,398 HQ SNVs (range: 72,867-77,373) called by both WES and WGS (**Fig. S3B**), 19,222 (18,823-20,024) of which were coding variants. The quality and distribution of the CD, GQ and MRR obtained with this combined calling process were similar to those previously reported for Unified Genotyper (**Fig. S4**).

We further investigated the HQ coding variants called exclusively by one method when the intersection of the two callers was used. We were able to separate the variants identified by only one technique into two categories: 1) those called by a single method and not at all by the other, which we refer to as fully exclusive variants, and 2) those called by both methods but filtered out by one method, which we refer to as partly exclusive variants. Of the HQ coding variants identified by WES only (105, on average, per sample), 61% were fully exclusive and 39% were partly exclusive. Of those identified by WGS only (692, on average) 21% were fully exclusive and 79% were partly exclusive. We performed Sanger sequencing on a random selection of 170 fully and partly exclusive WES/WGS variants. Of the 44 fully exclusive WES variants successfully Sanger sequenced, 40 (91%) were absent from the true sequence, indicating that most fully exclusive WES variants were false positives (**Table 2 and Dataset S1)**. By contrast, 39 (75%) of the 52 Sanger-sequenced fully exclusive WGS variants were found in the sequence, with the same genotype as predicted by WGS (including 2 homozygous), and 13 (25%) were false positives (**Table 2 and Dataset S1)**. These results are consistent with the observation that only 27.2% of the fully exclusive WES variants were reported in the 1000 genomes database (27), whereas most of the fully exclusive WGS variants (84.7%) were present in this database, with a broad distribution of minor allele frequencies (MAF) (**Fig. S5A**). Similar results were obtained for the partly exclusive variants. Only 10 (48%) of the 21 partly exclusive WES variants (including 3 homozygous) were real, whereas all (100%) of the 24 partly exclusive WGS variants (including 8 homozygous) were real.

**Table 2:**
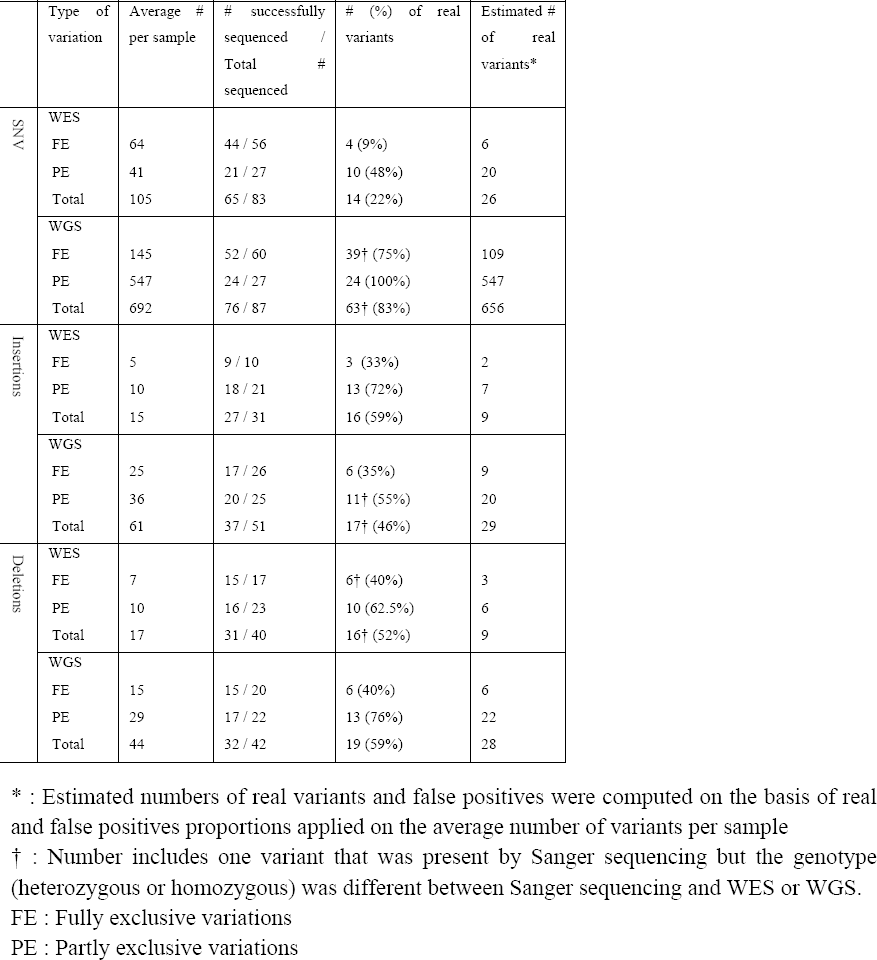
Results of Sanger sequencing for 170, 82 and 82 WES and WGS fully and partly exclusive SNVs, insertions and deletions respectively.

Using these findings, we estimated the overall numbers of false-positive and false-negative SNVs detected by these two techniques. WES identified a mean of 26 estimated real coding variants per sample (including 5 homozygous) that were missed by WGS, and a mean of 79 estimated false-positive variants. WGS identified a mean of 656 estimated real coding variants per sample (including 104 homozygous) that were missed by WES, and a mean of 36 estimated false-positive variants. We noted that most of the false-positive fully exclusive WGS SNVs were located in the three genes (ZNF717, OR8U1, and SLC25A5) providing the largest number of exclusive variants on WGS. Further investigations of the reads corresponding to these variants on the basis of blast (28) experiments strongly suggested that these reads had not been correctly mapped. Overall, we found that the majority of false-positive WGS fully exclusive variants (11/13) and only a minority of false-positive WES fully exclusive variants (4/40) could be explained by alignment and mapping mismatches. We then determined whether the exclusive WES/WGS SNVs were likely to be deleterious and affect the search for disease-causing lesions. The distribution of combined annotation-dependent depletion (CADD) scores (29) for these variants is shown in **Fig. S5B.** About 38.6% of the partly exclusive WES variants and 29.9% of the partly and fully exclusive WGS variants, which were mostly true positives, had a phred CADD score > 10 (i.e. they were among the 10% most deleterious substitutions possible in the human genome), and might include a potential disease-causing lesion. We also found that 54.6% of fully exclusive WES SNVs, most of which were false positives, had a phred CADD score > 10, and could lead to useless investigations. Overall, these results suggest that WGS is more accurate and efficient than WES for identifying true-positive SNVs in the exome.

We then compared WES and WGS for the detection of indels, following a strategy similar to that used for SNVs, and using Haplotype Caller, which is more appropriate than Unified Genotyper for indel detection (30). The mean number (and range) of insertions detected per sample was 5,795 (5,665-5,984) with WES and 6,443 (6,319-6,763) with WGS (**Fig. S3C**). On average, 5,313 insertions (85%) were called by both methods. The mean number (range) of deletions detected per sample was 7,530 (7,362-7,735) with WES and 6,259 (6,150-6,548) with WGS (**Fig. S3D**). On average, 5,383 deletions (74%) were detected by both methods. As for SNVs, the distributions of CD, GQ, and MRR for indels were of higher quality in WGS than in WES (**Fig. 1**). In particular, the distribution of CD was skewed to the right for both insertions and deletions. After applying the same filters as for SNVs (removing indels with CD < 8X, GQ < 20 and MRR < 0.2), we obtained a mean number of HQ insertions per sample of 4,104 (3,972-4,285) called by both WES and WGS (99.3% with the same genotype), 298 (248-413) called by WES only (5.3%), and 1,197 (974-1,400) called by WGS only (21.4%). We found that 4,121 HQ deletions (3,996-4,308) were called by both methods (99.5% with the same genotype), with the mean number of WES exclusive deletions (1,189; 1,015-1,419) similar to that of WGS exclusive deletions (1067; 871-1,215) (**Fig. S3**). We also investigated the HQ coding indels, which we defined as indels involving at least one bp included in a protein-coding exon. The mean number of HQ coding insertions per sample called by both WES and WGS was 247 (230-266). On average, 15 HQ coding insertions per sample were identified by WES only (33% of which were fully exclusive) and 61 were identified by WGS only (41% were fully exclusive). The mean number of HQ coding deletions per sample called by both WES and WGS was 240 (225-265). On average, 17 HQ coding deletions per sample were identified by WES only (41% of which were fully exclusive) and 44 were identified by WGS only (34% were fully exclusive).

The distribution of HQ indels by size is shown in **Fig. S6**. Most HQ insertions (57.1%) and deletions (57.3%) involved a single bp. We hypothesized that indels causing a frameshift in a coding region would be under stronger evolutionary constraints, and we investigated whether such coding regions were enriched in indels of a size corresponding to multiples of three bps. Consistent with this hypothesis, we observed that the number of insertions or deletions of a multiple of 3 bps was much higher for coding indels than for non-coding indels: >40% for coding indels and <12% for non-coding indels (**Fig. S6B and S6C**). These percentages were similar for coding indels called exclusively by WES or WGS (**Fig. S6B and S6C**), which suggests that most of these indels called exclusively by one method could be real. We Sanger sequenced a random selection of 164 coding indels exclusively called by WES or WGS. We found that the Sanger sequences of 32 of the 58 successfully sequenced WES-exclusive indels (55.2%) were consistent with WES findings (**Table 2 and Dataset S1**). Similarly, 36 (52.2%) of the 69 Sanger sequences obtained for WGS-exclusive indels were consistent with WGS findings (**Table 2 and Dataset S1**). These Sanger results indicate that, by contrast to what was found for SNVs, the estimated proportion of false-positives among exclusive HQ coding indels was equally high for both WES and WGS, at almost 50%. More indels were detected exclusively by WGS (61 + 44 = 105) than exclusively by WES (15 +17 = 32), so the number of real coding indels per sample detected by WGS and missed by WES was estimated to be higher (57) than that detected by WES and missed by WGS (18).

The results for indels should be interpreted bearing in mind that the Sanger sequencing of indels is more difficult than that of SNVs, for three main reasons. Indels can be complex, with a combination of insertion and deletion. The regions in which indels occur are often the hardest to sequence. For example, it is difficult to identify by Sanger sequencing a heterozygous deletion of 1 adenine (A) in a stretch of 20 consecutive A. And the analysis of the sequencing is harder, especially for heterozygous indels where it requires reconstructing manually the two alleles from long stretches of overlapping peaks. In this context, calls were difficult to make for a number of indels. We provide more detailed information about the analysis of the Sanger sequencing of indels in the methods. However, there are three arguments to suggest that our general findings for the WES/WGS comparison are valid. First, it was equally difficult to analyze Sanger sequences for WES-exclusive and WGS-exclusive indels. Second, the proportion of false positives was lower for partly exclusive than for fully exclusive indels for both WES (67.6% vs 37.5%) and WGS (64.8% vs 37.5%) (**Table 2**), as observed for SNVs. Finally, our Sanger results are consistent with the observed similar fractions of WES-exclusive and WGS-exclusive indels reported in the 1000 genomes database (**Fig. S5C**). Overall, these results indicate that the proportion of false-positive coding indels is similar for both WES and WGS.

The last step in our study was the comparison of WES and WGS for the detection of CNVs. The methods currently used to identify CNVs from WES data were already known to perform poorly for a number of technical reasons (31), including the fact that CNV breakpoints could often lie outside the regions targeted by the exome kit (32). A recent study comparing four WES-based CNV detection tools showed that none of the tools performed well and that they were less powerful than WGS on the same samples (16). We therefore restricted our analysis to the comparison of two classical WES-based methods, Conifer (33) and XHMM (32), with a well-known WGS-based method, Genome STRiP (34), for the detection of deletions in our six samples. As expected, more deletions were detected by WGS than by WES, with a total of 113 deletions (mean size: 23.7 kbs; range 0.8-182.5 kbs) detected over the six samples by Genome STRiP, 44 (mean: 45.6 kbs ; 0.3-644 kbs) detected by Conifer, and 30 (77.6 kbs; 0.1-2063 kbs) detected by XHMM. Ten of the 113 deletions detected by WGS (9%) were identified by Conifer, and 8 (7%) were detected by XHMM, including four detected by both Conifer and XHMM, consistent with the low concordance rates previously reported (16). We hypothesized that most true CNVs are common and should therefore be reported in public databases for CNVs, such as the Database of Genomic Variants (DGV) (35). The vast majority of deletions detected by WGS (105/113, 93%) were present in DGV, suggesting that most CNVs detected by WGS are true CNVs. Ten of the 14 deletions detected by WGS and at least one WES-based method (71.4%) were reported in DGV, whereas only 5/34 (14.7%) Conifer-exclusive and 5/22 (22.7%) XHMM-exclusive deletions were present in DGV. For 110 of the 113 deletions detected by WGS, one (24) or both (86) putative breakpoints were located outside the exome capture regions, providing a plausible explanation. Two of the three deletions with both breakpoints in the exome regions corresponded to the same 1.1 kb deletion identified in two different patients. This deletion was not identified by Conifer and XHMM, probably because only 3.4% of the 1.1 kb was covered by the exome kit. The third deletion was 29.1 kb long and was identified by both Conifer and XHHM. We also observed that most of the deletions (10/14) detected by WGS and at least one WES method belonged to the 20% of regions best covered by the exome kit. These results highlight the importance of both the size and the coverage ratio of the deletion by the exome kit, for optimal detection with the WES analysis methods currently available.

Finally, we investigated in more detail the coding regions and corresponding genes that were either poorly covered or not covered at all by the WES kit we used. We first determined, for each sample, the 1,000 genes with the lowest WES coverage. Up to 75.1% of these genes were common to at least four samples, and 38.4% were present in all six individuals. The percentage of exonic bps with more than 8X coverage for these 384 genes was, on average, 73.2% for WES (range: 0%-86.6%) and 99.5% for WGS (range: 63.6%-100%) (**Dataset S2A**). These genes with low WES coverage in all patients comprised 47 genes underlying Mendelian diseases, including three genes (*IMPDH1, RDH12, NMNAT1*) responsible for Leber congenital amaurosis, and two genes (*IFNGR2, IL12B*) responsible for Mendelian susceptibility to mycobacterial diseases (**Dataset S2A**). We then focused on the protein-coding exons that were fully outside the WES71+50 region **(Table 1)**. We restricted our analyses to the highest-quality protein-coding exons, those present in a consensus coding sequence (CCDS) transcript (21) with known start and end points of the coding sequence in the cDNA. These CCDS exons comprise a total of 46,227,845 bases which belong to translated regions, including 8,566,582 (18.5%) lying outside the WES71+50 region. The average CADD score for all possible variants was lower among the bases of the CCDS protein-coding exons not targeted by the kit (median= 7.362) as compared to the bases targeted by the exome kit (median= 14.87) **(Figure S5D)**. However, an important proportion (41.5%) of possible variants at bases not targeted by the kit had a CADD score >10 suggesting that potentially deleterious variations (with high CADD scores) might be missed by WES. We also found that 5,762 CCDS exons (3.1%), from 1,223 genes, were located entirely outside the WES71+50 region. Of these genes, 140 were associated with Mendelian diseases **(Dataset S2B).** We conducted the same analyses with the latest Agilent all-exon kit (v5+utr; 75 Mb), taking 50 bp flanking regions into account. We found that 2,879 CCDS exons (1.5%) were entirely excluded, and that these exons belonged to 588 genes, including 50 associated with monogenic disorders, such as *BRCA1,* the gene most frequently implicated in breast cancer. We also noted that, for these 2,879 CCDS exons, WGS detected a mean (range) of 436 (424-457) HQ SNVs and 16 (12-18) HQ indels.

## Discussion

Our findings confirm that WGS provides a much more uniform distribution of sequencing quality parameters (CD, GQ, MRR) than WES, as recently reported (14). The principal factors underlying the heterogeneous coverage of WES are probably related to the hybridization/capture and PCR amplification steps required for the preparation of sequencing libraries for WES (36). We also Sanger sequenced a large number of variations to obtain a high-resolution estimate of the number of false positives and false negatives obtained with WES and WGS (**Fig. 2**). All these analyses demonstrate that WGS can detect hundreds of potentially damaging coding SNVs per sample (∼3% of all HQ coding variants detected by WGS), about 16% of which are homozygous, including some in genes known to be involved in Mendelian diseases, that would have been missed by WES despite being located in the regions targeted by the exome kit (**Fig. 2**). The results are less clear-cut for indels, and should be interpreted more cautiously, as the current methods for identifying indels on the basis of both NGS and Sanger data are less reliable than those for SNVs. Our findings also confirm that WES is not currently a reliable approach to the identification of CNVs, due to the non contiguous nature of the captured exons, in particular, and the extension of most CNVs beyond the regions covered by the exome kit. In addition to the variants in the targeted regions missed by WES, a large number of exons from protein-coding genes, and non-coding RNA genes were not targeted by WES despite being fully sequenced by WGS (**Table 1**). It was noticeable that exons pertaining to CCDS transcripts were better covered than those that did not (**Table 1**). Finally, mutations outside protein-coding exons, or not in exons at all, might also affect the actual exome (the parts covered or not covered by WES), as mutations in the middle of long introns might impair the normal splicing of exons (37). These mutations would be missed by WES, but would be picked up by WGS (and selected as candidate mutations if the mRNAs were studied in parallel, for example by RNAseq).

**Fig. 2.**
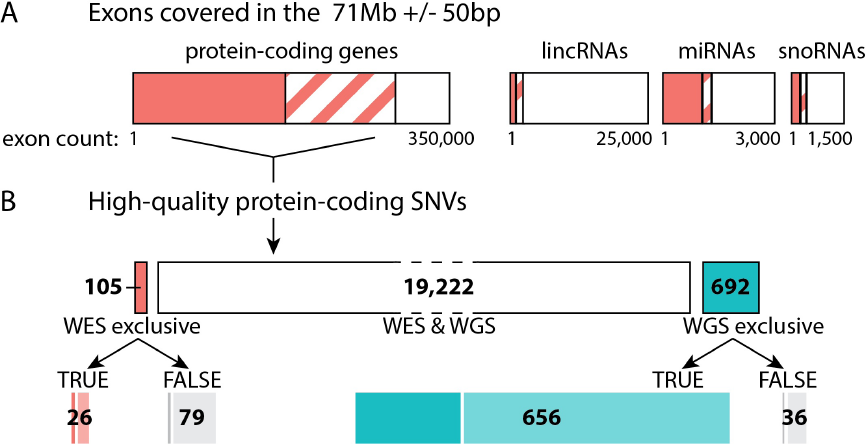
Diagram of the losses of single nucleotide variants (SNVs) at various levels associated with the use of WES. **(A)** Exons that were covered by the Agilent Sure Select Human All Exon kit 71Mb (V4 + UTR) with the 50 bp flanking regions. Exons fully covered are represented by boxes filled entirely in red; exons partly covered by boxes filled with red stripes; and exons not covered at all by white boxes. Numbers are shown in **Table 1**. Exons from protein-coding genes include exons encoding exclusively or partially UTRs, as well as exons mapping entirely to coding regions. **(B)** Number of high-quality coding SNVs called by WES and WGS (white box), by WES exclusively (red box), or by WGS exclusively (turquoise box). Details for the SNVs called exclusively by one method are provided below the figure. TRUE: estimate based on SNVs detected by Sanger sequencing. FALSE: estimate based on SNVs that were not detected by Sanger sequencing (**Table 2**). Darker boxes (red, gray, or turquoise) represent homozygous SNVs. Lighter boxes (red, gray, or turquoise) represent heterozygous SNVs.

Overall, our results show that WES and WGS perform very well for the detection of SNVs and indels, as more than 96.0% of HQ SNVs and 78.0% of HQ indels in the coding regions covered by the exome kit were called by both methods. The detailed analysis of the variants called exclusively by one approach showed that WGS was slightly but significantly more powerful than WES for detecting variants in the regions covered by the exome kit, particularly for SNVs. In addition, WGS is certainly more appropriate for detecting CNVs, as it covers all breakpoints and, of course, could detect variations in RNA- and protein-coding exome regions not covered by the exome kit. WGS currently costs two to three times as much as WES, but most of the cost of WGS (>90%) is directly related to sequencing, whereas WES cost is mainly due to the capture kit. Sequencing costs have greatly decreased and are expected to decrease faster than the cost of the capture kit. As an example, if sequencing costs were to decrease by 60% and capture kit costs remained stable, then the cost of WGS would approach that of WES. The cost of data analysis and storage should also be taken into account. In this rapidly changing economic context, specific cost/benefit studies are required and should take into account whether these NGS investigations are conducted for diagnostic or research purposes (15). These global studies should facilitate individual decisions, determining whether the better detection of SNVs, indels, and CNVs merit the additional cost of WGS, if WGS remains more expensive than WES. Finally, we carried out a detailed characterization of the variants and genes for which the two methods yielded the most strongly contrasting results, providing a useful resource for investigators trying to identify the most appropriate sequencing method for their research projects. We provide open access to all the scripts used to perform this analysis at the software website GITHUB (https://github.com/HGID/WES_vs_WGS). We hope that researchers will find these tools useful for analyses of data obtained by WES and WGS (4, 7, 9, 11, 38), two techniques that will continue to revolutionize human genetics and medicine in the foreseeable future.

## Materials and Methods

### Study subjects

The six subjects for this study (four female, two male) were collected in the context of a project on isolated congenital asplenia (39). They were all of Caucasian origin (two from USA, and one each from Spain, Poland, Croatia, and France), and unrelated. This study was overseen by the Rockefeller University IRB. Written consent was obtained from all patients included in this study.

### High-throughput sequencing

DNA was extracted from the Ficoll pellet of 10 mL of blood in heparin tubes. Unamplified, high-molecular weight, RNase-treated genomic DNA (4-6 µg) was used for WES and WGS. WES and WGS were performed at the New York Genome Center (NYGC) with an Illumina HiSeq 2000. WES was performed with Agilent 71Mb (V4 + UTR) single-sample capture. Sequencing was done with 2x100 base-pair (bp) paired-end reads, and 5 samples per lane were pooled. WGS was performed with the TruSeq DNA prep kit. Sequencing was carried out so as to obtain 30X coverage from 2x100 bp paired-end reads.

### Analysis of high-throughput sequencing data

We used the Genome Analysis Software Kit (GATK) best practice pipeline (17) to analyze our WES and WGS data, as detailed in the supporting information. We filtered out SNVs and indels with a CD < 8 or GQ < 20 or MRR < 20%, as previously suggested (24), with an in-house scrip. We used the Annovar tool (25) to annotate the resulting high-quality (HQ) variants. CNVs were detected in WES data with XHMM (32) and Conifer (33), and deletions were detected in WGS data with Genome STRiP (34), as detailed in the supporting information. All scripts are available from https://github.com/HGID/WES_vs_WGS.

### Analysis of Sanger sequencing

We randomly selected SNVs and indels detected exclusively by WES or WGS, for testing by Sanger sequencing. All the methods regarding the selection of variants, the design of primers, the sequencing of the variants, and the analysis of the Sanger sequences are provided in the supporting information.

## Acknowledgments

We would like to thank Vincent Barlogis, Carlos Rodriguez Gallego, Jadranka Kelecic, and Malgorzata Pac for the recruitment of patients, Fabienne Jabot-Hanin, Maya Chrabieh, and Yelena Nemirovskaya for their invaluable help, and the New York Genome Center for conducting WES and WGS. We thank the reviewers for their critical suggestions. Alexandre Bolze was funded by a fellowship from the Jane Coffin Childs Memorial Fund for Medical Research. The Laboratory of Human Genetics of Infectious Diseases is supported by grants from the March of Dimes (1-F12-440), National Center for Research Resources and the National Center for Advancing Sciences (NCATS) of the National Institutes of Health (8UL1TR000043), the St. Giles Foundation, the Rockefeller University, INSERM, and Paris Descartes University.

## Supporting information for methods

### Analysis of high-throughput sequencing data

We used the Genome Analysis Software Kit (GATK) best practice pipeline to analyse our WES and WGS data (1). Reads were aligned to the human reference genome (hg19) using the Maximum Exact Matches algorithm in Burrows-Wheeler Aligner (BWA) (2). Local realignment around indels was performed by the GATK (3). PCR duplicates were removed using Picard tools (http://picard.sourceforge.net). The GATK base quality score recalibrator was applied to correct sequencing artefacts. We called our 6 WES simultaneously together with 24 other WES using Unified Genotyper (UG) (3). as recommended by the software to increase the chance that the UG calls variants that are not well supported in individual samples rather than dismiss them as errors. All variants with a Phred-scaled SNP quality ≤ 30 were filtered out. The UG calling process in WGS was similar to that used for WES; we called our 6 WGS together with 20 other WGS. In both WES and WGS, the calling process targeted only regions covered by the WES 71 Mb kit + 50bp flanking each exon (4). When we expanded the WES regions with 100 and 200 bp flanking each exon as performed in some previous studies (5–7), we observed a higher genotype mismatch in variants called by WES and WGS, with a much lower quality of the WES variants located in those additional regions. Matched and mismatched genotype statistics, analyses of variant coverage depth (CD), i.e. the number of reads passing quality control used to calculate the genotype at a specific site in a specific sample, genotype quality (GQ), i.e. a phred-scaled value representing the confidence that the called genotype is the true genotype, and minor read ratio (MRR), i.e. the ratio of reads for the less covered allele (reference or variant allele) over the total number of reads covering the position where the variant was called, were performed using a homemade R software script (8).

We then filtered out variants with a CD < 8 or GQ < 20 or MRR < 20% a suggested in (9) using a homemade script.We used the Annovar tool (10) o annotate high quality (HQ) variants that were detected exclusively by one method. We checked manually some HQ coding variants detected exclusively by WES or WGS using the Integrative Genomics Viewer (IGV) (11), and we observed that some HQ coding WES exclusive variants, were also present in WGS but miscalled by the UG tool. To recall the UG miscalled SNVs, we used the GATK haplotype caller tool (HC) (3). Indels and SNVs were called simultaneously using HC on 6 WES and 6 WGS. We then split SNVs and indels into two combined vcf files. The same DP, GQ and MRR filters were applied for both SNV and Indels, and we used Annovar to annotate the HQ resulting variants.

CNVs were detected on WES data from our 6 samples together with 24 other samples originating from Europe using XHMM (12) and Conifer (13). For XHMM, we first calculated the depth of coverage in the 789,124 WES targets using GATK. XHMM was then run using default parameters to infer CNVs from read depths as previously described (12). For Conifer, the SVD-ZRPKM thresholds algorithm was used with the default parameters to find CNVs breakpoints (13). For WGS data, we ran Genome STRiP (14). ith the default parameters to detect large deletions on our 6 WGS together with 24 other WGS European samples from the 1000 genomes database (15). Genome STRiP looks for signatures of large deletions indicated by unusual spacing or orientation read pairs. We then kept only deletions that overlap with at least one WES targeted region. We looked whether the CNVs identified were present in the DGV database in February 2015 (16).

All scripts are available on https://github.com/HGID/WES_vs_WGS.

### Sanger sequencing methods

Selection of variants: We randomly selected variants detected exclusively by WES or WGS to test them by Sanger sequencing. We only sequenced once exclusive variants that were identified in multiple samples. We chose less variants in sample S1, as we had few gDNA available for this sample, and we could not test any of the variants in S2 because of absence of remaining gDNA. No other criteria (position, gene, CADD score, frequency, size of indel, etc.) was used for deciding which variants to Sanger sequence. For SNVs, we chose more variants in the two categories of WES fully-exclusive and WGS fully-exclusive as we first hypothesized (wrongly) that most, if not all, partly-exclusive variants would be real. The design and sequences of the primers will be provided on Figshare (www.figshare.com). Sanger sequencing was only attempted once for each variant.

Design of the primers: The first step was to create a bed file with each row representing a region of 400bp centered on the variants chosen for Sanger sequencing. The bed file was then uploaded in the UCSC genome browser using the ‘add custom tracks’ tab. The reference genome assembly used was GRCh37/hg19 (https://genome.ucsc.edu/cgi-bin/hgGateway). Fasta files with the sequence for each region were then downloaded from the UCSC website, and uploaded to BatchPrimer3 v1.0 (http://batchprimer3.bioinformatics.ucdavis.edu/cgi-bin/batchprimer3/batchprimer3.cgi) (17). We noticed that BatchPrimer3 worked better if the fasta files were copied and pasted rather uploaded using a link. We then requested for Sequencing primers using the following parameters: nb of return = 1 (1 towards 3’, and 1 towards 5’); sequencing start = -1; primer size: Min = 18, Opt = 22, Max = 25; primer Tm: Min = 55, Opt = 58, Max = 62; Max self complementarity = 8; Max 3’ self complementarity = 3. Lastly, variants for which one of the two primers was closer to 60bp to the variant were excluded from further sequencing and analysis. M13F or M13R sequences were added at the 5’-end of the forward or reverse primers. The full list of primers ordered is available in **Dataset S1**.

Sequencing of the variants: Amplification of the variants was performed using per reaction: H2O=11.5uL, 40% glycerol=4.5uL, 10X buffer (Denville without MgCl2)=2.25uL, MgCL2 (25mM)=0.9uL, dNTP (10mM)=0.225uL, primers (10uM)=0.5uL each, Taq Polymerase (Denville, #CB4050-2)=0.5uL, DNA=50-100ng. DNA was substituted by H2O in negative controls. 38 cycles of 94C (30’’), 60C (30’’), 72C (1’) were performed on a Veriti Thermal Cycler (Life Technologies). Sequencing PCR was done using the Big Dye 1.1 (Life Technologies) protocol with 1 uL of amplification PCR product and either the M13F or the M13R primer on a Veriti Thermal Cycler (Life Technologies). Lastly the samples were sequenced on a ABI 3730 XL sequencer (Life Technologies).

Analysis of the Sanger sequences: For SNVs, the analysis of the Sanger sequences was done using the DNASTAR SeqMan Pro software (v11.2.1) using the default settings. To facilitate the localization of the potential variants, we assembled the sequences obtained by Sanger with a 20bp fasta sequence centered on each variant. This sequence was obtained by creating a bed file of the region in the same way as described for the primer design. Variants where either the forward or reverse sequence did not work were excluded from the analysis and assigned a NA on the Sanger sequencing results **Dataset S1**. For indels, the Analysis of the Sanger sequences was much more difficult and it was not possible to use the DNASTAR SeqMan Pro software. Instead we used the software ApE (A plasmid Editor) to visualize every peak as clearly as possible. We then reconstructed the two alleles manually for each variant tested. For several indels, the analysis or results seemed intermediate. We considered that a variant was a false positive if: (i) there was no insertion or deletion at the place identified, or (ii) the size or sequence of the indel was incorrect, or (iii) the height of the peaks corresponding to the mutant allele were higher than the background noise usually observed - in practice we validated indels with sequencing peaks with a height >20% height of WT peaks. Lastly, we encountered several indels which were a combination of a deletion and an insertion. For example, the WT sequence would be: AAAAAAAAA and the mutated sequence would be: AAACAAA. Analysis of WES and WGS did not integrate these calls into one. We considered the results of WES or WGS true if WES or WGS called both the deletion of AAA and an insertion of a C in this example.

**Figure S1:**
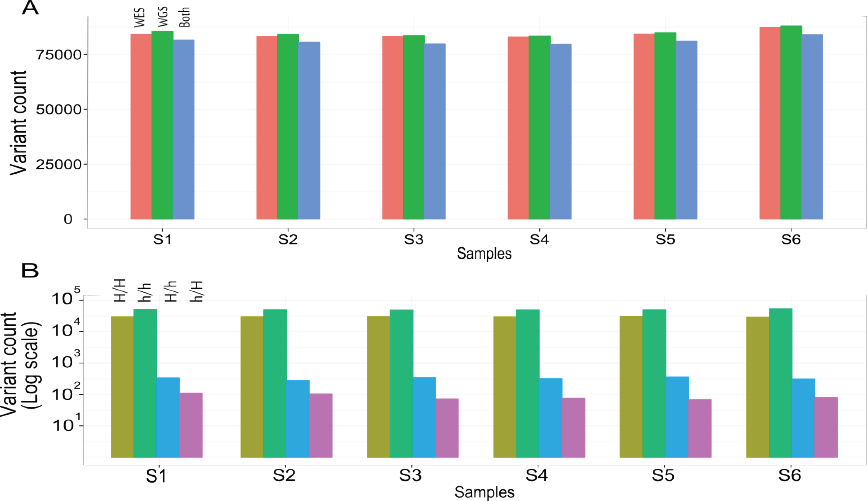
Number and general characteristics of single-nucleotide variants (SNVs) called by WES and WGS. **(A)** Total number of SNVs called by WES alone, WGS alone, and both platforms. **(B)** Characteristics of the SNVs called by both WES and WGS for each sample with four columns indicating the number of SNVs called homozygous by both methods (H/H, light green), called heterozygous by both methods (h/h, dark green), called homozygous by WES and heterozygous by WGS (H/h, blue), called heterozygous by WES and homozygous by WGS (h/H, purple)

**Figure S2:**
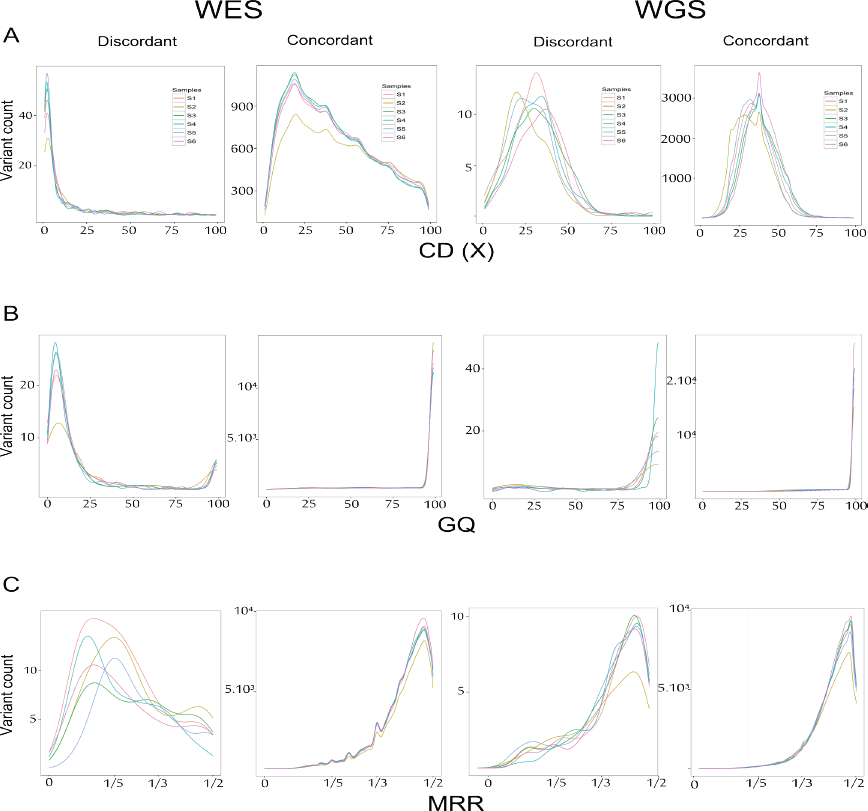
Distribution of the three main quality parameters for the SNVs with genotypes discordant between WES and WGS. **(A)** Coverage depth (CD), **(B)** genotype quality (GQ) score, and **(C)** minor read ratio (MRR). For each of the three parameters, four panels are shown: the two panels on the left show the characteristics of discordant and concordant SNVs in WES samples; the two panels on the right show the characteristics of discordant and concordant SNVs in WGS samples.

**Figure S3:**
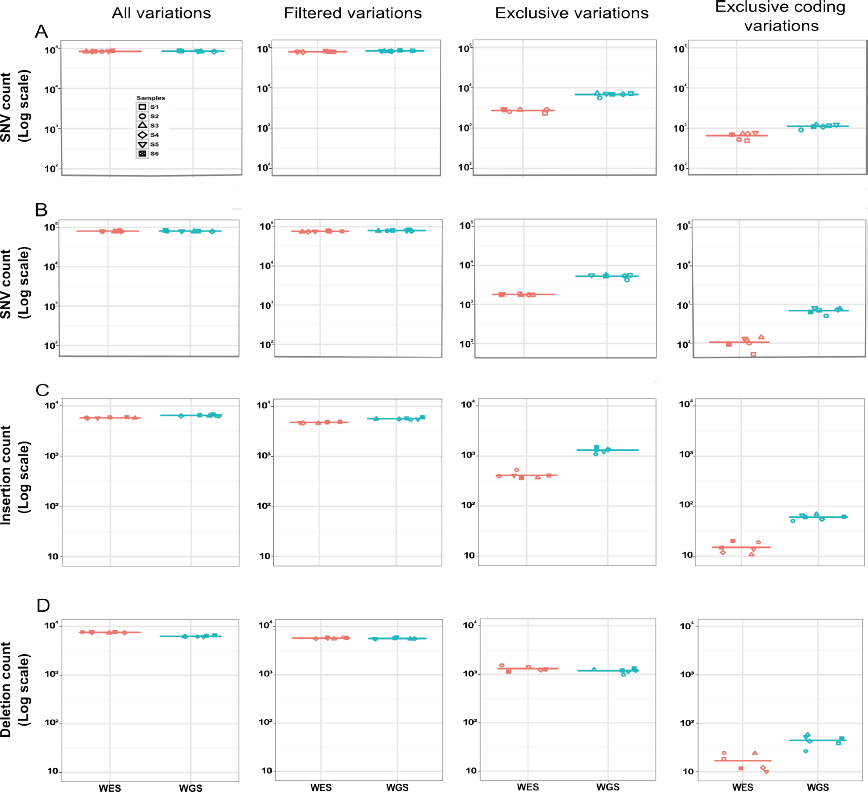
Numbers of variations in each WES or WGS sample following the application of various filters called with: **(A)** SNVs called using Unified Genotyper. **(B)** SNVs called using the intersection of Unified Genotyper and Haplotype Caller. **(C)** Insertions and **(D)** Deletions. Insertions and deletions were called using Haplotype Caller. For each of the four panels, we show from left to right: Total number of variations called by WES (red) or WGS (turquoise) for each sample; Total number of high-quality variations satisfying the filtering criteria: CD ȥ 8X, GQ ȥ 20 and MRR ȥ 0.2 called by WES (red) or WGS (turquoise) for each sample; Number of high-quality variations called by only one method, after filtering: high-quality exclusive WES variations (red) and high-quality exclusive WGS variations (turquoise); Number of exclusive WES (red) and exclusive WGS (turquoise) high-quality coding variations.

**Figure S4:**
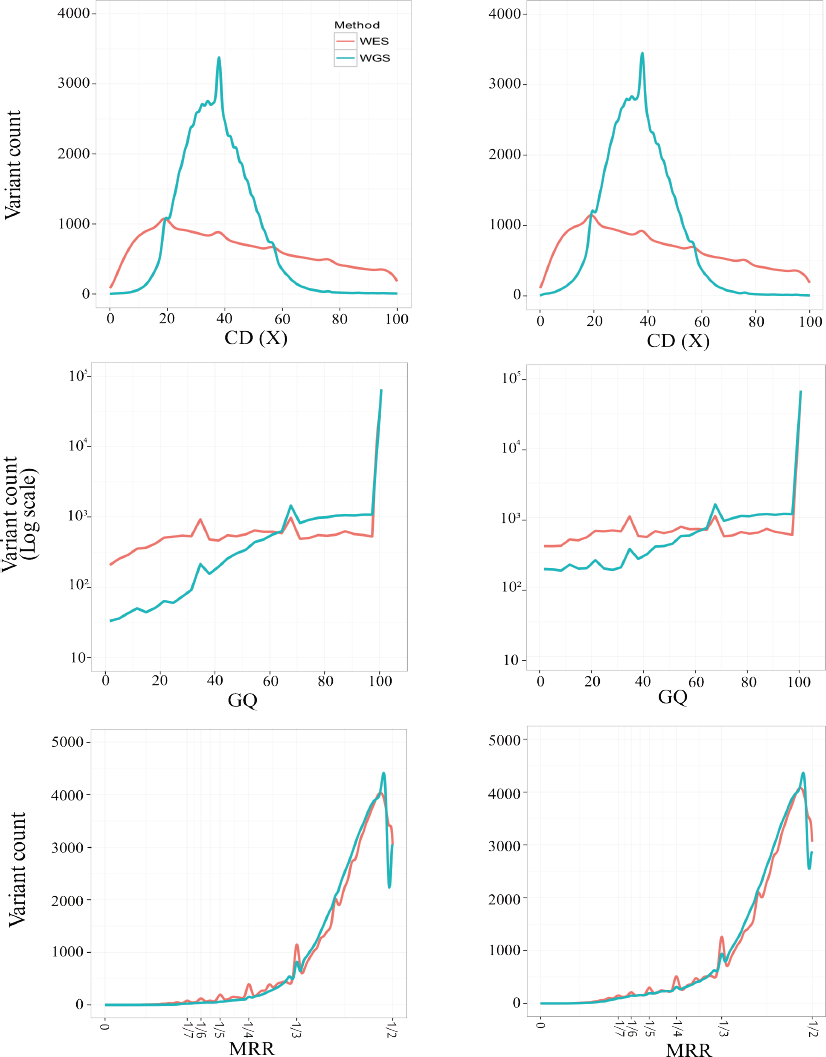
Comparison of the distribution of the three main quality parameters for the SNVs detected by WES or WGS, with either the intersection of Unified Genotyper and Haplotype Caller, or with Unified Genotyper alone. **(A)** Coverage depth (CD), **(B)** genotype quality (GQ) score, and **(C)** minor read ratio (MRR). For each of the three parameters we show: the average over the 6 WES (red) and the 6 WGS (turquoise) samples for the intersection of callers (left panel), and for Unified Genotyper alone (right panel)

**Figure S5:**
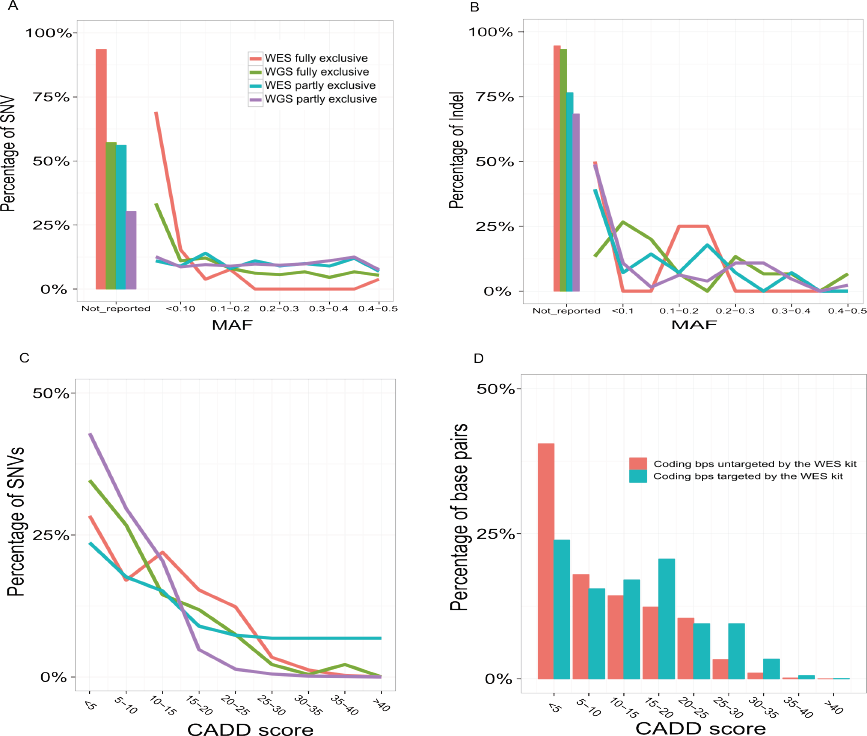
Characteristics of variations missed by WES or WGS. (A) and (B): Distribution of high-quality coding SNVs (A) and Indels (B) based on their presence and minor allele frequency (MAF) in the 1000 Genomes database. (C) and (D): Distribution of CADD (combined annotation-dependent depletion) scores (done on the version 1.2) for high quality coding SNVs identified exclusively by WES or by WGS (C); and for all base pairs included in the high-quality CCDS exons that were targeted (blue) or untargeted (red) with the 71Mb + 50Bp kit (D). For **(A)**, **(B)** and **(C)**: red represents fully exclusive high-quality WES coding variation never identified by WGS; turquoise represents partly exclusive high-quality WES coding variations identified by WGS but filtered out due to their poor quality; green represents fully exclusive high-quality WGS coding variations never called by WES; purple represents partly exclusive high-quality WGS coding variations identified by WES but filtered out due to their poor quality.

**Figure S6:**
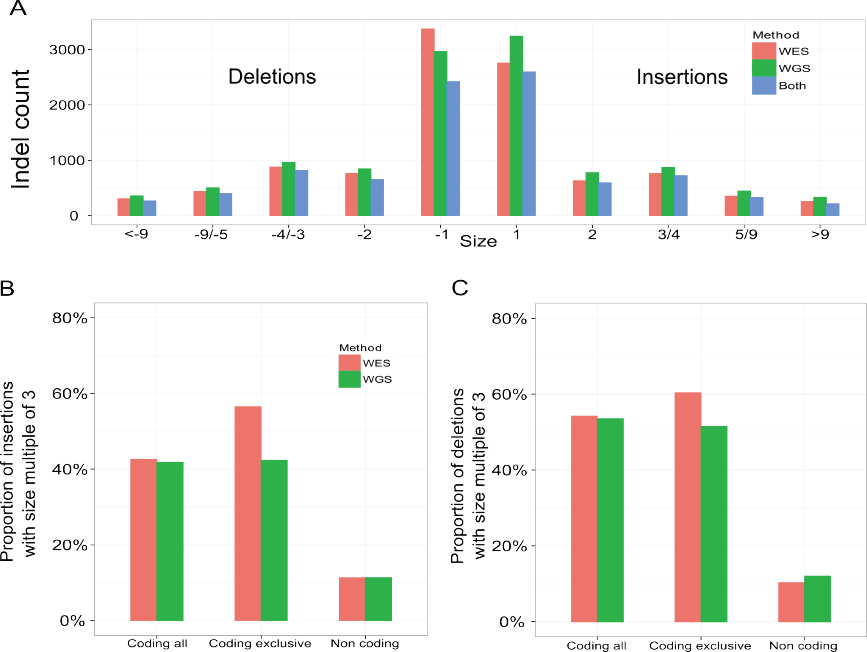
Distribution of high-quality Indels size: **(A)** Distribution of high quality Indels detected by WES (red), by WGS (green), and indicating those detected by both methods (blue) according to their size grouped in 5 categories : 1 bp, 2 bps, 3-4 bps, 5-9 bps and ȥ 10 bps; **(B)** Proportion of high-quality insertions with size multiple of 3 in coding and non-coding regions detected by WES (red) and WGS (green); **(C)** Proportion of high-quality deletions with size multiple of 3 in coding and non-coding regions detected by WES (red) and WGS (green). For coding regions we show both the total numbers of insertions/deletions and those that are WES or WGS exclusive.

**Table S1:** Reads and coverage statistics for each WES and each WGS.

| Sample | Total number of WES reads | Total number of WGS reads | Number of WES reads aligned in WES regions +/- 50 bps | Number of WGS reads aligned in WES regions +/- 50 bps | WES mean coverage in WES regions +/- 50 bps | WSG mean coverage in WES regions +/- 50 bps |
| --- | --- | --- | --- | --- | --- | --- |
| S1 | 98,792,738 | 1,370,493,918 | 64,696,895 | 34,737,193 | 72.1 | 38.7 |
| S2 | 124,483,242 | 1,303,868,290 | 80,970,674 | 31,743,245 | 90.3 | 35.3 |
| S3 | 86,822,862 | 1,477,715,120 | 57,970,027 | 37,322,280 | 64.5 | 41.5 |
| S4 | 89,521,104 | 1,438,287,290 | 59,084,117 | 36,600,011 | 65.9 | 40.7 |
| S5 | 98,002,162 | 1,301,586,284 | 62,673,065 | 33,102,614 | 69.9 | 36.8 |
| S6 | 100,056,600 | 1,445,702,068 | 68,002,983 | 37,619,386 | 75.8 | 41.9 |
| Mean | 99,613,118 | 1,389,608,828 | 65,566,294 | 35,187,455 | 73.1 | 39.2 |

